# Variant pathogenic prediction models VSRFM and VSRFM-s, the importance of splicing and allele frequency

**DOI:** 10.1101/430975

**Authors:** JL Cabrera-Alarcon, J Garcia-Martinez

**Author notes:** Both authors are corresponding author.

## Abstract

Currently, there are available several tools to predict the effect of variants, with the aim of classify variants in neutral or pathogenic. In this study, we propose a new model trained over ensemble scores with two particularities, first we consider minor frequency allele from gnomAD and second, we split variants based on their splicing for training each specific model. Variants Stacked Random Forest Model (VSRFM) was constructed for variants not involved in splicing and Variants Stacked Random Forest Model for splicing (VSRFM-s) was trained for variants affected by splicing. Comparing these scores with their constituent scores used as features, our models showed the best outcomes. These results were confirmed using an independent data set from Clinvar database, with similar results.

## BACKGROUND

High-throughput next-generation genomic technology is a disruptive event in the research of genetic based diseases. However, it implies a challenge in discerning pathogenic from benign variants. For this purpose, researchers have built different scope developed algorithms. Up to date, there are several tools for predicting variants pathogenicity, including ensemble scores, that gather information from different single pathogenic predictors, improving the overall outcome in variants classification.

On the other hand, conservation scores give evolutionary measures over a specific position, and therefore variant consequence. Nevertheless, the conservation degree depends on the considered phylogenetic group.

Alternative splicing (AS) is a major biological mechanism for rising protein diversity in organisms. Complexity in AS is correlated with evolution and tissue complexity (1). Thus, conservation level may vary drastically depending on considered phylogenetic groups and in a higher rate than it does in exonic variants that directly modify protein. Because of this particular behavior it could be interesting to split variants depending on splice events participation.

Here, we present two stacked meta learners for deleteriousness classification, trained in two different data sets: Variants Stacked Random Forest Model (VSRFM), trained in variants not involved in splicing and Variants Stacked Random Forest Model for splicing (VSRFM-s), trained in variants involved in splicing events.

## MATERIAL AND METHODS

### Study models

We trained two random forest models, VSRFM for not splice variants data set and VSRFM-s in splice involved variants data sets, using randomForest v-2.5.1 R package (2).

### Datasets

To build VSRFM and VSRFM-s we used supervised machine learning over a composed data set which contains pathogenic and benign variants, obtained selecting unique variants from five benchmark datasets *HumVar (3), ExoVar (4), VariBench(5), predictSNP (6) and SwissVar (7)*. Variants were annotated using Variant Effect Predictor web interface for GRCh37, with metaLR, metaSVM, REVEL, DANN, phastCons and phyloP scores from dbNSFP v3.5a, ada score and rf score, Condel, CADD and gnomAD allele frequency. Study data set contained 82922 variants, which were split in two different datasets, one with 2145 splice implicated variants (1322 pathogenic and 823 neutral variants) and another with 80777 not splice implicated variants (38878 pathogenic and 41899 neutral variants).

We tested the accuracy of VSRFM and VSRFM-s against their constitutive scores in a set of 10664 variants selected from Clinvar archive, classified as benign or pathogenic variants, not presented in our training dataset, annotated with all needed scores (8). These variants were split again in two data sets according to their participation in splicing, in 10109 not splice Clinvar variants (6965 pathogenic and 3144 neutral variants) and 555 splice Clinvar variants (481 pathogenic and 74 neutral variants).

### Features

For models building we used 12 different scores for not splice variants data set, 6 general esemble functional predictor scores, MetaLR (9), MetaSVM (9), REVEL (10), DANN (11), CADD (12) and Condel (13), four conservation scores, phastCons (14) and phyloP (15) conservation score based on the multiple alignments of 100 vertebrate and 20 mammalians genomes and gnomAD allele frequencies for exome variants (16). Variants without minor allele frequency (MAF) value in gnomAD database are considered rare variants and automatically assigned minimum MAF value 0.0000081. For splice variants we considered the addition of two ensemble scores ada score an rf score, for altered splicing prediction from dbscSVN (17). Missing values for splice variants and not splice variants dataset are represented in table 1s. For both data sets variant data imputation was carried out using randomForest v-2.5.1 R package (2).

For testing feature correlation, we used spearman correlation test, represented using ggplot2 v-2.2.1 R package (18).

### Receiver operating characteristic (ROC) curves

Receiver operating characteristic (ROC) curves were used to compare deleteriousness classification ability between our scores and their different components scores considered in this study. ROC curves plots and its areas under the curve (AUC) were made using ROCR v-1.0-7, cvAUC v-1.1.0 and pROC v-1.12.1 R packages (19–21). For both models, AUC was computed as an average of AUC obtained in 10-fold cross validation. For ROC curve comparison we used pROC v-1.12.1.

## RESULTS

The correlation study between features in not splice variants showed that MetaLR, MetaSVM and REVEL had very strong correlation level (Spearman correlation coefficient r_s_ > 0.8) also seen between CADD and DANN and between Condel and CADD, figure 1s. It is interesting to notice that while there is very strong correlation between phastCons and phyloP in vertebrates, that correlation is moderate in mammalians (Spearman correlation coefficient 0.39 > r_s_ > 0.60). The rest of possible comparisons between functional ensemble scores are strong (Spearman correlation coefficient 0.59 > r_s_ > 0.79), figure 1s. In training splice variants decreases the overall correlation level between features. Both dbscSVN scores showed a very strong correlation level, figure 2s.

**Figure 1.**
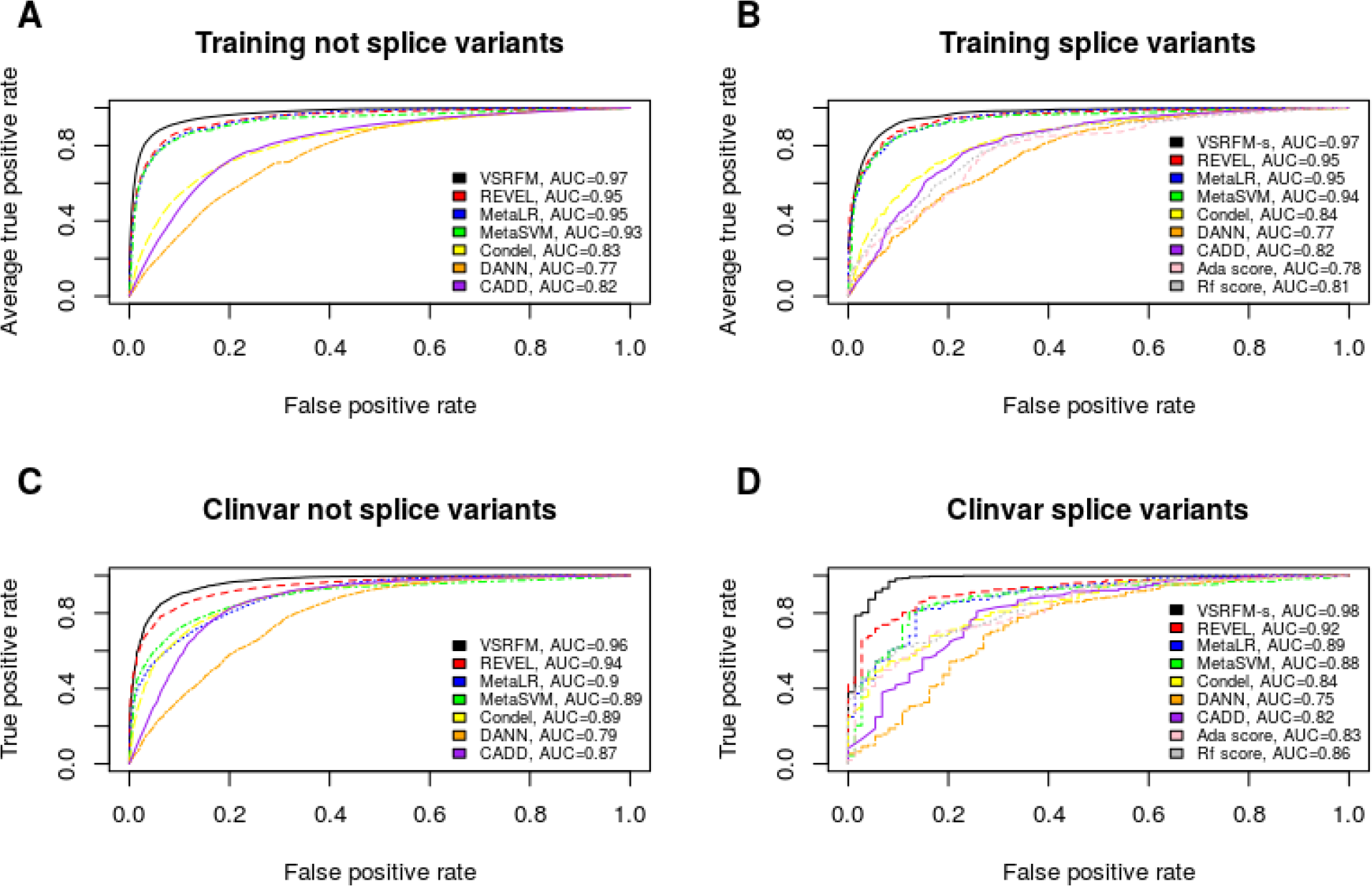
ROC curves for models in training not splice variant data (A), training splice variant data (B), Clinvar not splice variant data (C) and Clinvar splice variant data (D).

ROC curves for our two models, VSRFM, VSRFM-s, and their constituent predictor scores both for splice and not splice non-synonym exomic variants, are shown in figure 1. VSRFM and VSRFM-s and their component scores, using area under the ROC curve were statistically significant better in both in non-synonym splice involved variants as in not splice involved variants, p-value < 1*10^−06^, both in our training data set (AUC = 0.97 in splice and in not splice variants), as in Clinvar selected variants (AUC = 0.97 in not splice variants and AUC = 0.98 in splice variants), table 2s.

According to our data, we purpose a cutoff for VSRFM = 0.4599167 (0.9987139 sensibility and 0.9900236 specificity) and for VSRFM-s = 0.5079833 (0.9992436 sensibility and 0.9963548 specificity), figure 2.

**Figure 2.**
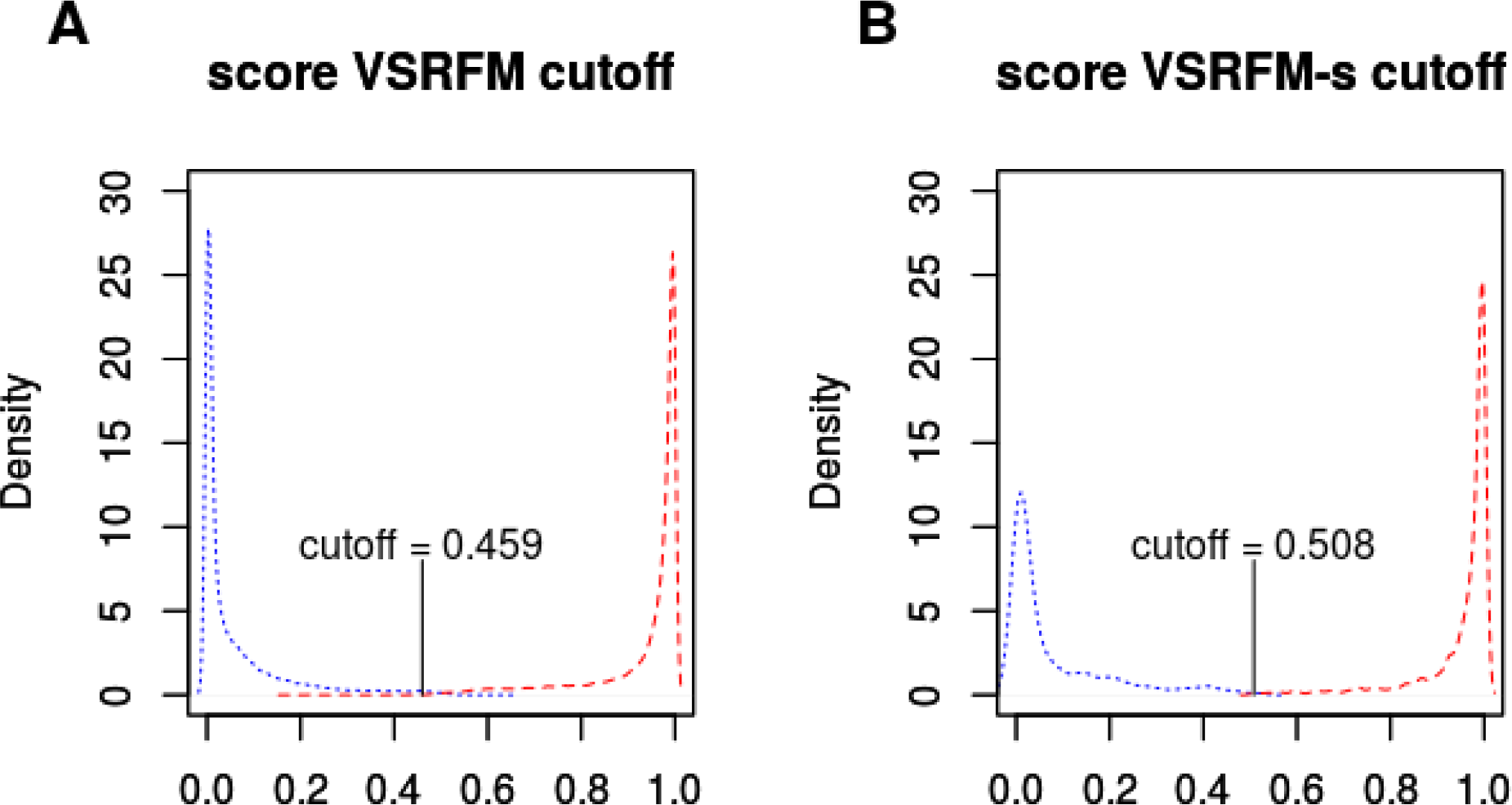
VSRFM and VSRFM-s scores. VSRFM score in deleterious (red) and neutral (blue) variants and purposed cutoff (A). Distribution of VSRFM-s score and chosen cutoff value (B).

Relative importance of VSRFM and VSRFM-s, as decrease in Gini impurity for features in VSRFM and VSRFM-s are represented in table 1. The most important constituent score in both classifiers were REVEL. Looking at these results gnomAD MAF information plays a relevant role in outcome weighting for both models. In not splice variants, though scores conservation information are included in other features, as REVEL, were included in random forest training, given that they still provided additional information, however they have very low relevance in splice variants.

**Tabla 1.** Feature relative importance.

| **Tabla 1. Feature relative importance.** |  |  |
| --- | --- | --- |
|  | **Decrease Gini impurity** |  |
|  | **VSRFM** | **VSRFM-splicing** |
| **MetaLR** | 3852.26 | 90.40 |
| **MetaSVM** | 3104.22 | 68.72 |
| **REVEL** | 5479.67 | 137.11 |
| **gnomAD** | 2633.51 | 78.90 |
| **DANN** | 637.41 | 10.71 |
| **CADD** | 980.03 | 19.88 |
| **Condel** | 974.15 | 23.78 |
| **phastCons100way_vertibrate** | 209.61 | 2.81 |
| **phastCons20way_mammalian** | 375.30 | 6.99 |
| **phyloP100way_vertibrate** | 831.56 | 11.55 |
| **phyloP20way_mammalian** | 411.38 | 7.80 |
| **Ada score** | - | 14.64 |
| **Rf score** | - | 19.43 |

## DISCUSSION

We have developed two scores to discriminate neutral from probably deleterious variants, VSRFM-s and VSRFM in splice and not splice involved variants, respectively.

According to the level of correlation observed between features, the high correlation level showed may be due to the strong correlation between these features and our target/classification group. This observation is especially strong between REVEL, metaLR and metaSVM which could be consequence of sharing lots of single scores, despite they are different algorithms. This could mislead random forest to give more importance to a feature over other highly correlated, as REVEL, that presented the highest Gini impurity decrease, and hence had the highest importance over some other classifiers as metaLR or metaSVM. But this is not an issue that leads to overfit our random forest model, as could be seen in overall performance in Clinvar data.

Regarding to the ROC curve comparison, the results obtained in this study were consistent with Sieh and coworkers, about the relative overall performance of used scores (10). In the comparison between VSRFM and VSRFM-s and their constituent functional scores, our models showed the largest discriminative power, followed by REVEL, metaLR and metaSVM, both in training data set and in Clinvar data set, for their specific subset of variant type (splice/not splice). Unlike the strategy adopted by other authors, focusing in MAF to select working variants, we decided to include this information to train the random forest algorithm. In this way, in accordance with Gini impurity decrease, gnomAD allele frequency presented high relevance in weighting the final outcome.

The reduced relevance of conservation scores in VSRFM-s development, points to the fact that in spite of there are several conserved positions involved in splicing, this conservation depends on considered. In this way these different levels of conservation were not recovered by the two conservation levels considered in this study.

For all exposed, we can conclude that VSRFM and VSRFM-s are tools that improve pathogenic mutation detection.

